# Epidemiology of bacterial contamination of inert hospital surfaces and equipment in critical and non-critical care units: a Brazilian multicenter study

**DOI:** 10.1101/793034

**Authors:** Dayane Otero Rodrigues, Laís da Paixão Peixoto, Erica Tatiane Mourão Barros, Julianne Rodrigues Guimarães, Bruna Clemente Gontijo, Jaisa Leite Almeida, Lucas Guimarães de Azevedo, Júlia Cristina Oliveira e Lima, Deyse Silva Câmara

**Affiliations:** Centro de Ciências Biológicas e da Saúde, Universidade Federal do Oeste da Bahia, Barreiras, Bahia, Brasil; Laboratório de Microbiologia, Curso de Biomedicina, Centro Universitário Luterano de Palmas, Palmas, Tocantins, Brasil; Curso de Medicina, Centro de Ciências Biológicas e da Saúde, Universidade Federal do Oeste da Bahia, Barreiras, BA, Brasil

## Abstract

The hospital environment is an important reservoir of microorganisms, including multidrug-resistant pathogens, which can cause in-patient contamination and healthcare-related infections. The objective of this study was to analyze the epidemiology of bacterial contamination (contaminated sites, pathogen species and their antimicrobial susceptibility, and tracking of multidrug-resistant microorganisms - MDR) of inert hospital surfaces and medical equipment in two public hospitals in Northern Brazil. This was a cross-sectional study with 243 samples (*n* = 208, from Hospital A; and *n* = 35, from Hospital B) collected by friction with swabs moistened in Brain Heart Infusion from inert surfaces and equipment. The samples were cultivated and bacterial species were identified by the classical approach and tested for their susceptibility through agar diffusion assay according to the Clinical and Laboratory Standards Institute (CLSI). Most inert surfaces and equipment analyzed presented bacterial contamination (95.5%). *Staphylococcus aureus* was the main pathogen of clinical significance detected both in Hospital A (61.8%) and B (68.6%). Hospital A showed higher rates of isolated MDR bacteria than Hospital B, especially in the Adult Intensive Care Unit, which included methicillin-resistant *Staphylococcus aureus* (MRSA) (52.7%), Enterobacteria resistant to 4^th^ generation cephalosporins (19.4%), and multidrug-resistant *Pseudomonas aeruginosa* (2.78%). The failures in the prevention and control of infections in the two hospitals analyzed reinforce the need for a revised protocol for cleaning and disinfection of inert surfaces and medical equipment, and for regulation of antibiotic dispensing, mainly in the AICU of Hospital A, which was found to be a reservoir of MDR pathogens. This study is innovative because it is the pioneer in Western Bahia that describes the epidemiology of contamination of hospital surfaces, opportuning futures studies in this field.

## Introduction

Healthcare-associated infections (HAI) are a major public health concern commonly associated with extended length of hospital stay. HAI account for high hospital costs and contribute to increased morbidity and mortality of infected patients [1].

HAI are usually caused by pathogenic bacteria that may emerge from the patient’s endogenous microflora during antibiotic therapy in approximately 70% of the cases [2,3]. HAI may also be acquired from the exogenous environment (30% of the cases) in that the hospital setting plays a significant role in contagion and transmission outbreaks [3,4].

In the hospital setting, patients, staff and visitors represent the main reservoir of microorganisms, whereas secondary reservoirs include all environments where nutrients, moisture, and temperature are suitable for microbial survival, such as air humidifiers and nebulizers [5,6]. In addition, dry and inanimate surfaces can also serve as a reservoir of pathogens [3,5,7-9], as in mattresses and bed frames [4,10,11], door knobs [11,12], and even in medical equipment such as stethoscopes and ultrasound devices [7-11]. Contamination of these surfaces contributes to pathogen spreading and, as a result, development of horizontal infections [13-14].

Overall, surfaces can be directly contaminated by bacteria from colonized and/or infected patients or from the hands of health professionals [5,7]. Highly touched surfaces (e.g. bed frames, stethoscopes, bedside tables, and door knobs) [11-13] may be contaminated by common bacteria of the hand microbiota. More importantly, MDR bacteria have been detected in medical equipment and contact surfaces, especially in critical care units [10,15,16].

Studies carried out in Brazil and North America have reported contamination of hospital surfaces by bacteria resistant to antibiotics, especially methicillin-resistant *S. aureus* (MRSA) [4,17,18] and vancomycin-resistant enterococci [19,20]. These findings indicate that patients and staff are at risk of contamination by pathogens associated with high mortality rates against which treatment options are restricted.

Given the importance of in-hospital transmission, HAI studies have looked into the epidemiology of hospital-related bacterial contamination to propose preventive measures to reduce the contamination and dissemination of resistant pathogens. This study aimed to analyze the epidemiology of bacterial contamination (contaminated sites, pathogen species and their antimicrobial susceptibility, and tracking of multidrug-resistant microorganisms - MDR) of inert hospital surfaces and medical equipment in two public hospitals in Northern Brazil.

## Materials and methods

This was a multicenter, cross-sectional study carried out in the *Hospital Geral Público de Palmas* (HGPP) [Palmas Public General Hospital] (A) located in the state of Tocantins, Brazil, and in the *Hospital Municipal Eurico Dutra* [Eurico Dutra Municipal Hospital] (B) located in the city of Barreiras, Western Bahia, Brazil.

Hospital A is a large teaching hospital which provides tertiary care and is a reference center for medium and high complexity healthcare assistance. The hospital contains approximately 400 beds and is a major health center for the state of Tocantins and neighboring states in Brazil. It includes two intensive care units (ICUs) – a pediatric unit (PICU) with 3 rooms and 8 beds, and an adult unit (AICU) with 26 beds distributed among 18 rooms; pharmacy; laboratory; operating room (OR); elective and emergency surgery services; emergency room (ER) with three on-call specialties – orthopedics, internal medicine, and surgery; conventional hospital ward, and home care and outpatient facilities, with an average 3,500 appointments *per* month.

Hospital B is a medium-sized and medium complexity general hospital which provides services such as hospital admission, diagnostic and therapeutic support services, emergency and outpatient care. Patients are admitted through spontaneous and referred demand from the city of Barreiras and surrounding region. The hospital has a total of 10 beds in the Internal Medicine Ward (IMW), 29 beds in the Surgical Ward (SW) and 4 OR with 5 post-anesthesia recovery beds in the SW.

### Sample collection

A total of 243 samples (*n* = 208, Hospital A; *n* = 35, Hospital B) were collected from the following surfaces and medical equipment: Hospital A - ER (door knobs, *n* = 56); SW (door knobs, *n* = 20); AICU (heart monitors, *n* = 18; infusion pumps, *n* = 18; medication tables, *n* = 18; side bed frames, *n* = 18; with a total *n* = 72); and PICU (side bed frames, *n* = 16; bed headboards, *n* = 8; bed frame feet, *n* = 8; mattresses, *n* = 8; bedside tables, *n* = 8; stethoscopes, *n* = 8; samples from the sinks, *n* = 4; with a total *n* = 60); Hospital B - IMW (bed headboard, *n* = 3, mattresses, n = 2; side bed frame, *n* = 2; saline stand, *n* = 1; door knobs, *n* = 3, with a total *n* = 11); SW (bed headboard, *n* = 5, mattresses, *n* = 3; side bed frame, *n* = 1; saline stand, *n* = 3; countertops, *n* = 1; door knobs, *n* = 4; with a total *n* = 17); OR (surgical light, *n* = 4; stretcher, *n* = 3; with a total *n* = 7). The selected hospital surfaces included those highly touched or nearby patients.

The samples were collected in the morning and afternoon from July to October 2018 (Hospital A) and April 2018 (Hospital B) by friction with sterile swabs moistened in broth Brain Heart Infusion (BHI, Oxoid, Basingstoke, Hampshire, England) from selected surfaces. The swabs were placed immediately into sterile tubes and transported in insulated boxes to the *Centro Universitário Luterano de Palmas* (CEULP) [Palmas Lutheran University Center] (Hospital A) and to the *Universidade Federal do Oeste da Bahia* (UFOB) [Federal University of Oeste da Bahia] (Hospital B) and incubated at 37 °C for 24 h.

### Bacterial culture

After incubation, the samples were vortexed and subcultured onto Mannitol Salt Agar, MacConkey, and Pseudomonas agar supplemented with blood to isolate *S. aureus*, coagulase-negative staphylococci (CoNS), and Gram-negative bacteria (Enterobacteriaceae family, non-fermenters), respectively.

Next, clinically significant bacterial species [21] were identified by the classical approach as follows:

- *S. aureus* and CoNS: fermentation of Mannitol Salt Agar, Gram staining, catalase test, coagulase test, DNase Agar;
- Enterobacteriaceae family: growth on MacConkey agar, cytochrome oxidase, Gram staining, lactose fermentation (Triple Sugar Iron Agar -TSI), biochemical tests (B Enterokit Probac of Brazil);
- Non-fermenter Bacilli (*Pseudomonas aeruginosa*): growth on Pseudomonas agar, odor, colony morphology, cytochrome oxidase, growth at 42°C, Gram staining, oxidation in Hugh Leifson medium;

The bacterial strains were then tested for their antimicrobial susceptibility in vitro by the agar diffusion technique as recommended by the Clinical and Laboratory Standards Institute [22].

Bacterial strains were considered as multidrug-resistant (MDR) if showing resistance to at least three classes of antimicrobials [23,24] and associated phenotypes. Some examples included MRSA, Enterobacteria resistant to 4^th^ generation cephalosporins and quinolone-resistant *P. aeruginosa*, according to definitions of the European Center for Disease Prevention and Control [23].

## Results

Bacterial contamination was detected in 94.7% of the 208 sampled surfaces and equipment in Hospital A. A total of 233 bacterial isolates were obtained therefrom, with no microbial growth in only 11 surfaces and equipment in the PICU. Moreover, 100% of the 35 sampled surfaces and equipment in Hospital B showed bacterial contamination.

The microbiological analysis of the surfaces in Hospital A showed a predominance of common bacteria of the human flora, such as *S. aureus* (62.0%) and nosocomial bacteria, such as Enterobacteria (24.0%) and *P. aeruginosa* (7%). In Hospital B, there was a predominance of environmental microorganisms, such as Gram-positive Bacilli (41%) and *Micrococcus* spp. (9.0%) (Figure 1).

**Figure 1.**
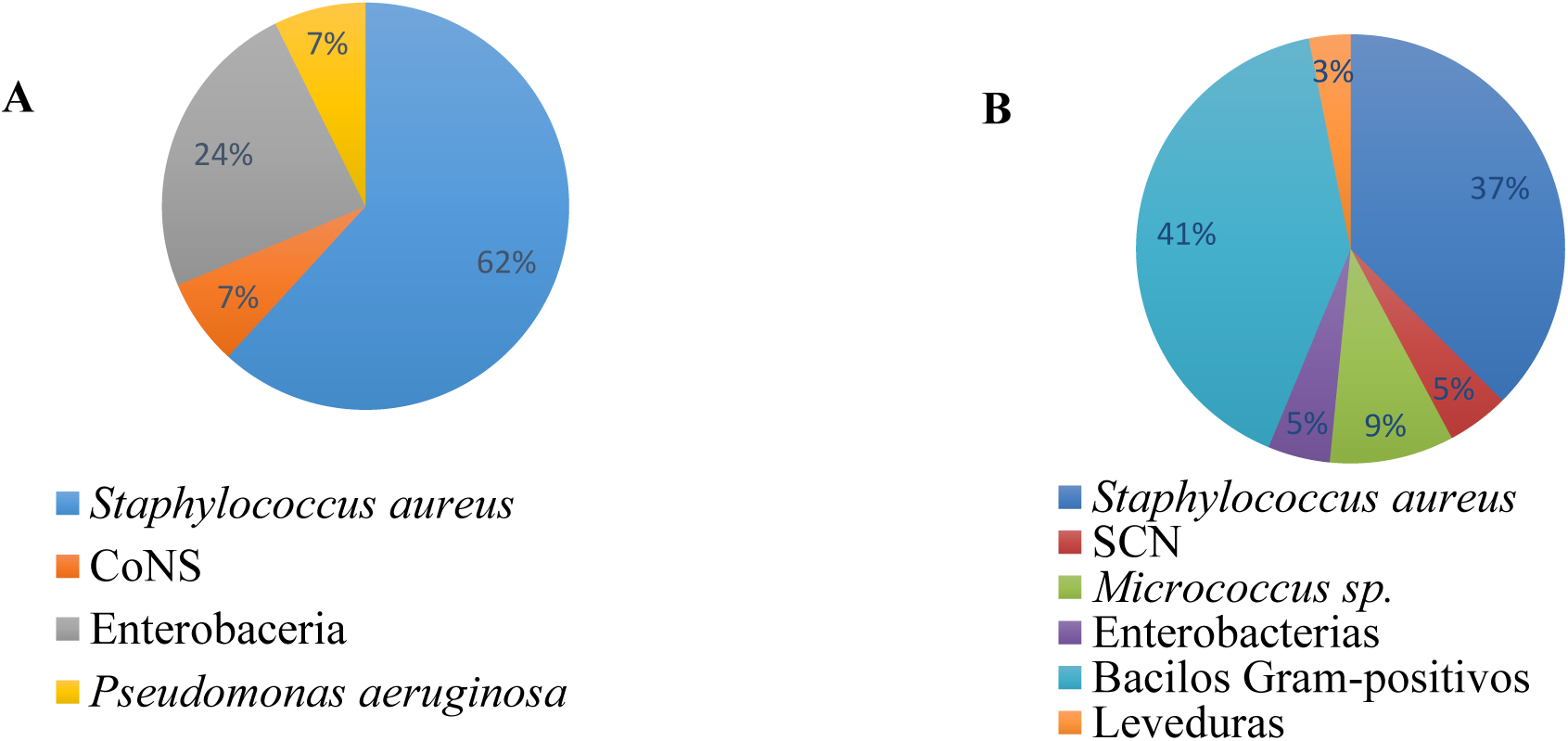
Frequency of microorganisms recovered from inert surfaces and medical equipment in the Hospital Geral Público de Palmas-TO, Brazil (A) and Hospital Municipal Eurico Dutra in the city of Barreiras, BA, Brazil (B).

In Hospital A, the ER door knobs were mostly contaminated with *S. aureus* (53.3%), but also with enteric bacteria (30.4%), with a high frequency of samples contaminated with *P. aeruginosa* (64.7%) (Table 1). The AICU showed the highest frequency of *S. aureus* colonization, particularly on heart rate monitors (83.3%). Infusion pumps and side bed frames were mostly contaminated with *P. aeruginosa* (16.7%). The PICU was frequently contaminated not only with *S. aureus* in all stethoscopes analyzed, but also with CoNS. As shown in Table 1, 10 out of the 16 CoNS samples found throughout the study originated from the PICU.

**Table 1.** Frequency of clinically important microorganisms isolated from inert surfaces and medical equipment in the Hospital Geral Público de Palmas -TO, Brazil (A).

| Hospital units | Surfaces (N) | *S.aureus*<br>N (%) | CoNS<br>N (%) | Enterobacteria<br>N (%) | *P. aeruginosa*<br>N (%) | Total of bacteria (N) |
| --- | --- | --- | --- | --- | --- | --- |
| ER | door knobs (56) | 49 (53,3) | 4 (4,3) | 28 (30,4) | 11 (11,9) | 92 |
| OR | door knobs (20) | 12 (60,0) | 2 (10,0) | 5 (25,0) | 1 (5,0) | 20 |
| AICU | heart monitors (18) | 15 (83,3) | - | 3 (16,7) | - | 18 |
|  | infusion pumps (18) | 13 (72,2) | - | 3 (16,7) | 2 (11,1) | 18 |
|  | medication tables (18) | 14 (77,8) | - | 4 (22,2) | - | 18 |
|  | side bed frames (18) | 11 (61,1) | - | 6 (33,3) | 1 (5,6) | 18 |
| PICU | side bed frames (16) | 8 (57,1) | 1 (7,4) | 5 (35,7) | - | 14 |
|  | bed headboards (8) | 3 (50,0) | 2 (33,3) | 1 (16,7) | - | 6 |
|  | bed frame feet (8) | 3 (50,0) | 2 (33,3) | - | 1 (16,7) | 6 |
|  | mattresses (8) | 3 (75,0) | 1 (25,0) | - | - | 4 |
|  | bedside tables (8) | 3 (42,8) | 3 (42,8) | 1 (14,3) | - | 7 |
|  | stethoscopes (8) | 8 (100,0) | - | - | - | 8 |
|  | samples from the sinks (4) | 2 (50,0) | 1 (25,0) | - | 1 (25,0) | 4 |
| **Total (N)** | **208** | **144 (61,8)** | **16 (6,9)** | **56 (24,0)** | **17 (7,3)** | **233** |

N: total sample size; %: Percentage; ER: Emergency Room; OR: Operating Room; AICU:
Adult Intensive Care Unit; PICU: Pediatric Intensive Care Unit; CoNS: Coagulase-Negative Staphylococci.

The surfaces of Hospital B were contaminated with *S. aureus* in the following sites: bed headboards (66.7%) in the IMW, saline stands (100.0%) in the SW and surgical lights (100.0%) in the OR (Table 2).

**Table 2.**
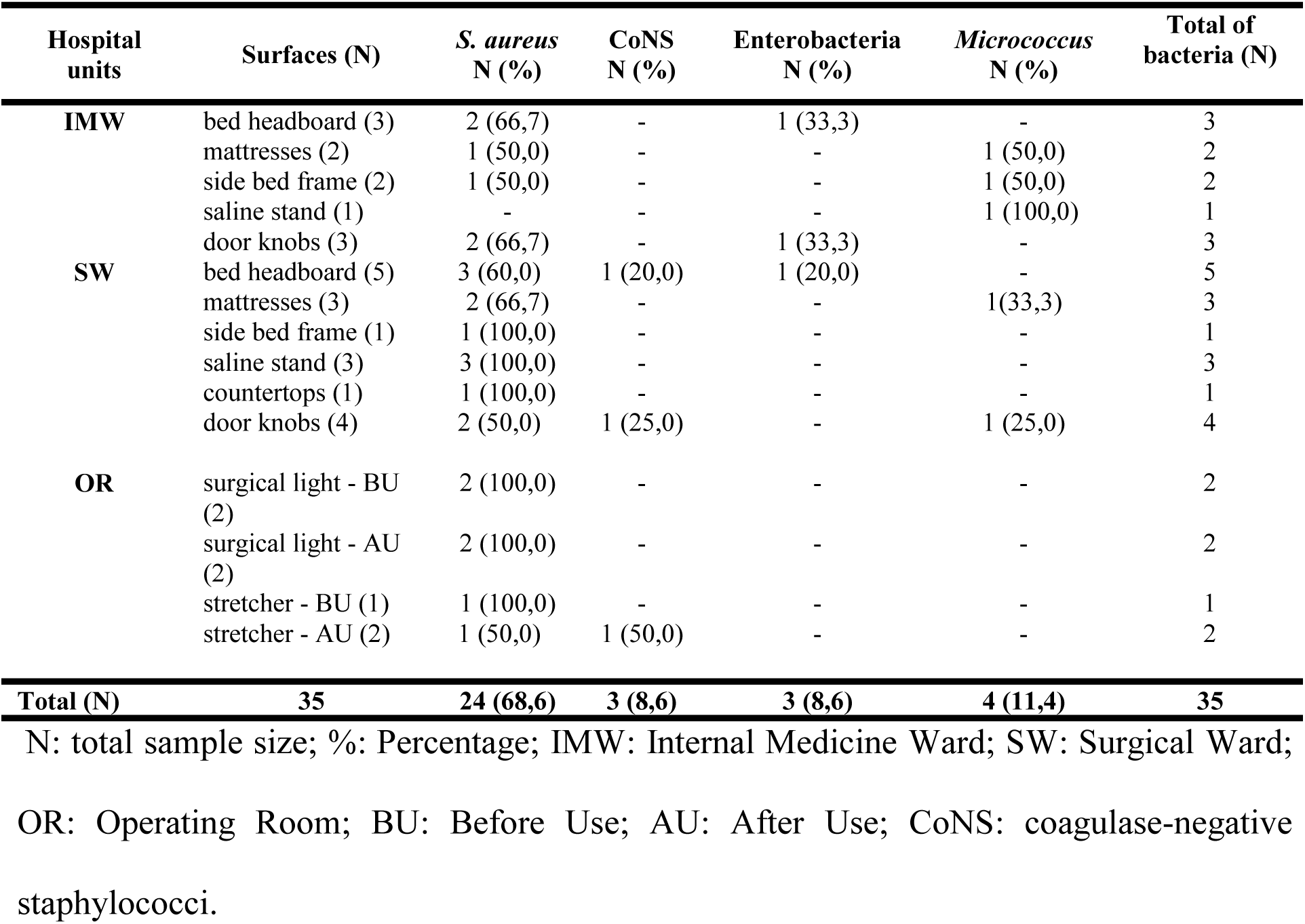
Frequency of clinically important microorganisms isolated from inert surfaces and medical equipment in the Hospital Municipal Eurico Dutra in the city of Barreiras, BA, Brazil (B).

| Hospital units | Surfaces (N) | *S. aureus* N (%) | CoNS N (%) | Enterobacteria N (%) | *Micrococcus* N (%) | Total of bacteria (N) |
| --- | --- | --- | --- | --- | --- | --- |
| **IMW** | bed headboard (3) | 2 (66,7) | - | 1 (33,3) | - | 3 |
|  | mattresses (2) | 1 (50,0) | - | - | 1 (50,0) | 2 |
|  | side bed frame (2) | 1 (50,0) | - | - | 1 (50,0) | 2 |
|  | saline stand (1) | - | - | - | 1 (100,0) | 1 |
|  | door knobs (3) | 2 (66,7) | - | 1 (33,3) | - | 3 |
| **SW** | bed headboard (5) | 3 (60,0) | 1 (20,0) | 1 (20,0) | - | 5 |
|  | mattresses (3) | 2 (66,7) | - | - | 1(33,3) | 3 |
|  | side bed frame (1) | 1 (100,0) | - | - | - | 1 |
|  | saline stand (3) | 3 (100,0) | - | - | - | 3 |
|  | countertops (1) | 1 (100,0) | - | - | - | 1 |
|  | door knobs (4) | 2 (50,0) | 1 (25,0) | - | 1 (25,0) | 4 |
| **OR** | surgical light - BU (2) | 2 (100,0) | - | - | - | 2 |
|  | surgical light - AU (2) | 2 (100,0) | - | - | - | 2 |
|  | stretcher - BU (1) | 1 (100,0) | - | - | - | 1 |
|  | stretcher - AU (2) | 1 (50,0) | 1 (50,0) | - | - | 2 |
| **Total (N)** | **35** | **24 (68,6)** | **3 (8,6)** | **3 (8,6)** | **4 (11,4)** | **35** |

N: total sample size; %: Percentage; IMW: Internal Medicine Ward; SW: Surgical Ward;
OR: Operating Room; BU: Before Use; AU: After Use; CoNS: coagulase-negative staphylococci.

Hospital A showed higher rates of MDR bacteria recovered from surfaces than Hospital B, especially in the AICU, which included MRSA (52.7%), Enterobacteria resistant to 4^th^ generation cephalosporins (19.4%) and multi-resistant *P. aeruginosa* (2.78%). The OR in both Hospital A and B had the lowest frequency of MDR bacteria, as shown in Table 3.

**Table 3.**
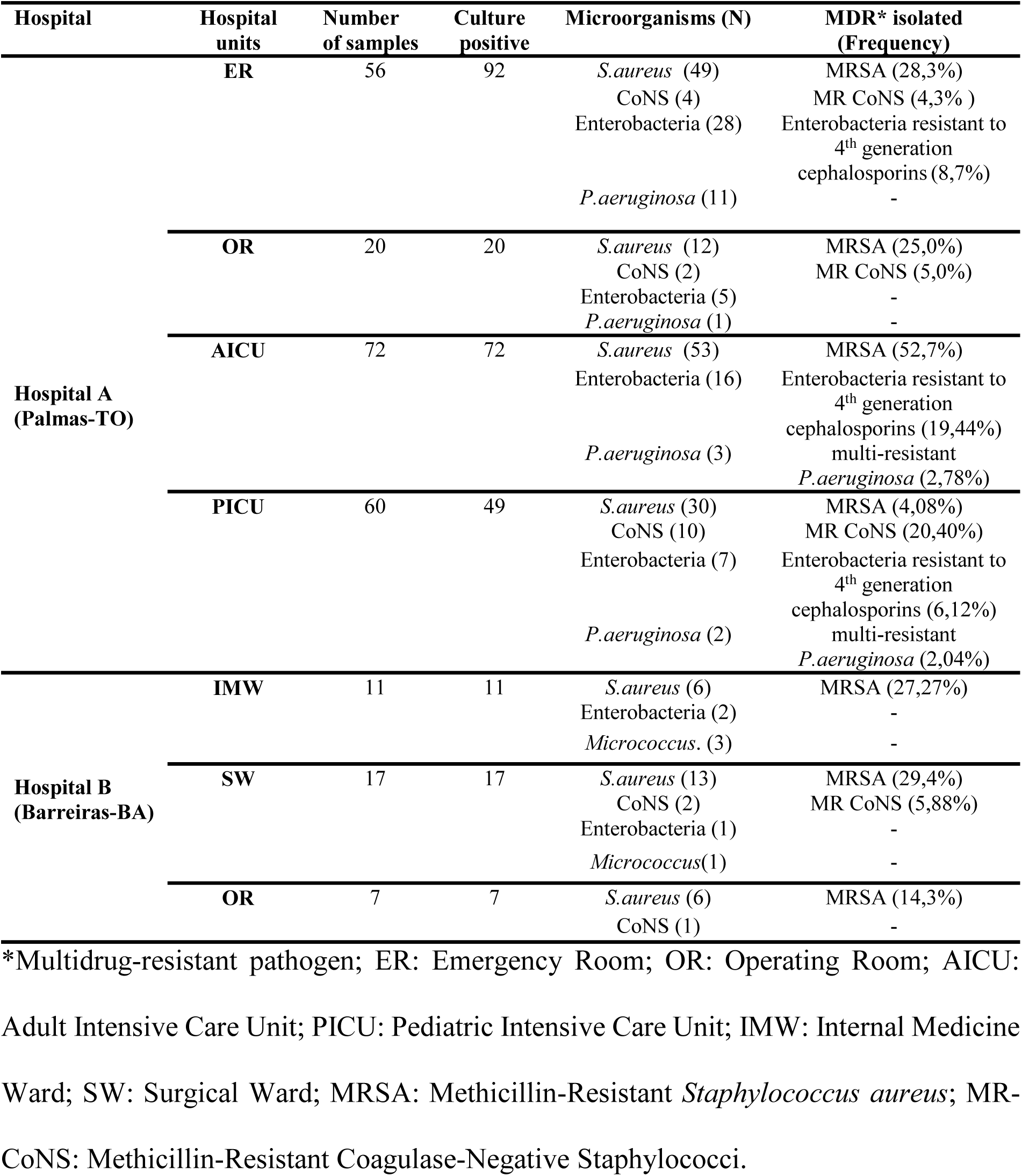
Multi-drug resistant microorganisms recovered from inert surfaces and equipment of hospitals located in Northern Brazil.

## Discussion

Inert hospital surfaces and medical equipment can be a reservoir of MDR pathogens. Understanding the epidemiology of bacterial contamination in this setting is essential to prevent in-patient contamination and HAI development.

Bacterial pathogens can survive and remain viable on inert surfaces and equipment due to their ability to form biofilms and to environmental factors such as surface porosity and humidity [7,25]. This ability to adapt to environmental stress works as a major factor driving pathogen thriving and dissemination. Our results are consistent with these premises as 94.7% (Hospital A) and 100.0% (Hospital B) of the inanimate surfaces analyzed (*n* = 243) in general hospitals located in Northern Brazil were contaminated with bacterial pathogens. Other factors should also be considered, which include proximity of colonized and/or infected patients to the surfaces analyzed; the physical structure of the hospital; the adoption of antibiotic administration programs; as well as issues with the cleaning and disinfection protocols of surfaces and medical equipment. These findings corroborate those of national surveillance studies carried out in ICUs in Center-Western [26] and Northern Brazil [15]. Consistent with this, international studies [2,5,27] have shown that only less than 50% of hospital surfaces are properly cleaned and disinfected with germicides [28-30]. These alarming findings strongly suggest that the hospital environment can act as a reservoir of pathogens and enable their cross-transmission to the patient.

While bacteria, viruses, and fungi can commonly thrive in environmental reservoirs [21,31,32], the etiology of microbial contamination in our study was specific to each hospital. Hospital A had surfaces and equipment contaminated with human pathogens, whereas Hospital B was contaminated with environmental bacterial contaminants. These differences may be explained by several factors [19,33] such as hospital size and cleaning/disinfecting regimens. Hospital A is a large hospital providing medium and high complexity healthcare assistance, with an ICU demanding appropriate cleaning/disinfection. In addition, admitted patients have an increased length of hospital stay and are often prescribed antibiotics indiscriminately. A combination of these circumstances may lead to selection of nosocomial pathogens such as MRSA, *P. aeruginosa* and Enterobacteria, which ultimately contaminate the surrounding environment. On the other hand, Hospital B is a medium-sized hospital providing medium complexity healthcare assistance, which does not present all the aggravating factors described for Hospital A. In Hospital B, admitted patients have a better prognosis and the presence of common bacterial contaminants on hospital surfaces (e.g., Gram-positive Bacilli and *Micrococcus* spp.), which are widely distributed in the environment, is justified [21]. The bacterial contamination present therein may result from the patient’s transient skin microbiota, even though opportunistic infections may be developed in immunocompromised individuals [21].

*S. aureus* was the main microorganism recovered from the surfaces and equipment of both Hospital A (61.8%) and B (68.6%). In Hospital A, there was a predominance of *S. aureus* in door knobs in the ER (53.3%), heart monitors (83.3%) and medication tables (77.8%) in the AICU, and stethoscopes (100.0%) and mattresses (75.0%) in the PICU. In Hospital B, *S. aureus* was the main pathogen recovered from door knobs and bed headboards (66.7%), saline stands (100.0%) and mattresses (66.7%). These percentages are similar to those reported in the national literature [4] for mattresses (72.0%) and higher than international percentages [34] for beds (34%) and door knobs (26.0%). Our data show that highly touched surfaces are prone to be contaminated with *S. aureus*, a microorganism commonly found in the humans’ hand microbiota. This suggests that the hands of health professionals were the main vector of contamination of surfaces in the units analyzed, although this was not investigated. Surfaces and equipment can be contaminated and, if not are properly disinfected, re-contaminate sanitized hands during the interruption of patient care to touch the surfaces and equipment such as keyboards, tables, door knobs, stethoscopes etc. The surfaces become permanently contaminated by pathogens [5,7,19] and the hospital environment turns into a reservoir of highly transmissible microorganisms.

In our study, bedside tables of the PITU were found to be contaminated with CoNS (42.8%), although at a frequency lower than that of PICU clinicians’ mobile phones in Kuwait (62.9%) [35]. CoNS are main cause of late-onset sepsis in developed countries^36^ and early neonatal infections in Canadá [36], USA [37] and Brazil [38]. Despite the fact that CoNS are not considered pathogenic under normal circumstances, their presence on objects with frequent manual contact (e.g., bedside tables and areas close to patients such as bed headboards and bed frame feet) may pose a risk of contamination and development of infection in hospitalized children. This fact requires attention by local health teams and indicates a need for revision of surface cleaning/disinfection protocols at the PICU in Hospital A.

Our findings showed a high percentage of methicillin-resistant isolates of CoNS and mostly *S. aureus*. The frequency of MRSA recovered from the ER (28.3%) was similar to that reported in the literature [12] (20.0%), while the frequency of MRSA recovered from the AICU (52.7%) was higher than that found in Brazil [39] (41.8%) and lower than international rates [40] (67.3%). The presence of MRSA in both hospitals analyzed reflects the persistence of this pathogen in the environment [34], as observed in another tertiary hospital in the Middle East in five rooms and two nursing stations in the AICU and PICU. MRSA strains often exhibit multidrug resistance, including resistance to beta-lactam antibiotics, and exhibit strong virulence as a result of a multiplicity of factors acting simultaneously to evade the host’s defenses. Some of which include the production of enzymes and toxins, intracellular invasion and proliferation, dissemination into tissues and organs, and ability to form biofilms [4,27,41-43]. These virulence factors mediate bacterial adhesion to inert hospital surfaces and medical equipment, which may explain the high prevalence rates of MRSA found in this study, although biofilm production was not investigated herein.

The contamination of surfaces and equipment with Gram-negative bacilli is not as studied as the contamination with MRSA, VRE and *C. difficile* [3]. A study [44] reported the presence of MDR Enterobacteria in 22.2% of 18 sampled surfaces, which is consistent with our findings (19.44%) for the AICU of Hospital A. European data [45] showed a high level of contamination by *P. aeruginosa* in sink isolates and tap biofilms. In line with this, our study recovered *P. aeruginosa* from 25% of sink samples, which attests that this pathogen is commonly associated with humid sites in the hospital environment and is the main microorganism isolated in ICUs [46]. In contrast, from a total of seventeen *P. aeruginosa* isolates found in this study, eleven were non-MDR isolates from door knobs of the ER, which is not a humid environment. This may be associated with the presence of *P. aeruginosa-*infected patients in beds near contaminated door knobs. Once more, this reinforces the understanding that health professionals’ hands work as a vector of pathogen transmission. In our study, MDR *P. aeruginosa* isolates were recovered at a low frequency from AICU (2.78%) and PICU (2.04%) facilities, which differs from the literature showing 40.8% [47] prevalence of MDR *P. aeruginosa* in two hospitals in the West region of Santa Catarina state in Brazil. Even so, our results confirm the imminent risk of contamination of critically ill patients.

In our study, we observed that the AICU of Hospital A was highly contaminated with MDR pathogens. This is in line with observational studies that recovered MDR microorganisms from critical care units [5,27,40]. The environmental contamination of ICUs has been a worrisome issue, since severely ill patients are prone to develop infections. It is fundamental to adopt standard measures to prevent and control hospital-acquired infection, considering the patients’ risk of death, the proximity of the beds and the presence of monitoring and support equipment, which are highly touched and susceptible to contamination. It is also worth noting the widespread use of antibiotics in critical units, which select MDR clones both in the patient and in the environment, in addition to the fact that viable MDR bacteria have been isolated from biofilm surfaces and furniture after terminal cleaning of ICUs [5,48]. Evidence has suggested that ICUs are epicenters of MDR microorganisms, which was confirmed in our study.

## Conclusions

There was a high frequency of contaminated surfaces and equipment in our study, with isolation of difficult-to-treat MDR phenotypes, e.g. MRSA, MR-CoNS, Enterobacteria resistant to 4^th^ generation cephalosporins and MDR *P. aeruginosa*. These findings raise concern and point to the need to review the protocols of prevention and control of infections in the hospitals analyzed, mainly Hospital A. The cleaning/sanitizing of inert surfaces and equipment and antibiotic dispensing procedures should also be looked at in critical units, since antibiotics can act as a selective force of resistant bacterial strains. Routine and terminal disinfection of environmental surfaces and medical equipment with germicides is recommended by the literature [32] to decrease the frequency and level of contamination, especially highly touched surfaces near the patient. There is a need for greater attention into hand hygiene before and after contact with patients as much as with the areas close to them. These actions together could reduce contamination of hospital surfaces and equipment and the possibility of cross-transmission of pathogens while minimizing the risk of contamination of patients and the development of HAI.

Taken altogether, the association between environmental contamination and the epidemiology of HAI is complex. This study is the pioneer in Western Bahia that describes the epidemiology of contamination of hospital surfaces, opportuning futures studies in this field. Further research should better understand the correlation of bacterial contamination with the underlying pathology of hospitalized patients close to contaminated surfaces and use molecular biology tools to determine the similarity of strains recovered from surfaces and patients, in order to track the circulation and origin of MDR pathogens during cross-infection. Our results showed a high frequency of contaminated surfaces and equipment, and critical care units acting as reservoir of MDR pathogens, including MRSA, MR-CoNS, Enterobacteria resistant to 4^th^ generation cephalosporins and MDR *P. aeruginosa*, and these pathogens are commonly associated with poor prognosis, difficult-to-treat HAI. These results may assist health professionals of the analyzed hospitals, especially Hospital A, in the adoption of preventive and control measures for HAI and reveal the need for reflection on the indiscriminate use of antimicrobials.

## Acknowledgements

We would like to thank to board directors from *Hospital Geral Público de Palmas* (HGPP) [Palmas Public General Hospital], the board directors from *Hospital Municipal Eurico Dutra* [Eurico Dutra Municipal Hospital] for their institutional consent.

## References

1. Bonnet V, Dupont H, Glorion S, Aupée M, Kipnis E, Gérard JL, et al. Influence of bacterial resistance on mortality in intensive care units: a registry study from 2000 to 2013 (IICU Study). J Hosp Infect. 2019; 102 (3): 317–24.

2. Weber DJ, Rutala WA, Miller MB, Huslage K, Sickbert-Bennett E. Role of hospital surfaces in the transmission of emerging health care-associated pathogens: Norovirus, Clostridium difficile, and Acinetobacter species. Am J Infect Control. 2010; 38 (5 Suppl 1): S25–33.

3. Weber DJ, Rutala WA. Understanding and Preventing Transmission of Healthcare-Associated Pathogens Due to the Contaminated Hospital Environment. Infect Control Hosp Epidemiol. 2013; 34 (5): 449–52.

4. Ferreira AM, Andrade D, Almeida MTG, Cunha KC, Rigotti MA. Colchões do tipo caixa de ovo: um reservatório de *Staphylococcus aureus* resistente à meticilina? Rev Esc Enferm. 2011; 45: 161–6.

5. Russotto V, Cortegiani A, Raineri SM, Giarratano A. Bacterial contamination of inanimate surfaces and equipment in the intensive care unit. J Intensive Care. 2015; 3:54–61.

6. Trindade RC, Bonfim ACR, Resende MA. Conjuntivais flora microbiana de pessoas clinicamente normais que trabalham em um ambiente hospitalar. Braz J Microbiol. 2000; 31 (1): 12–6.

7. Russotto V, Cortegiani A, Fasciana T, Iozzo P, Raineri SM, Gregoretti C, et al. What Healthcare Workers Should Know about Environmental Bacterial Contamination in the Intensive Care Unit. Biomed Res Int. 2017; 2017: 6905450.

8. Boyce JM. Environmental contamination makes an important contribution to hospital infection. J Hosp Infect. 2007; 65 (Suppl 2): 50–4.

9. Rossi D, Devienne KF, Raddi MSG. Influência de fluídos biológicos na sobrevivência de *Staphylococcus aureus* sobre diferentes superfícies secas. Rev Ciênc Farm Básica Apl. 2008; 29: 211–4.

10. Adams CE, Smith J, Watson V, Robertson C, Dancer SJ. Examining the association between surface bioburden and frequently touched sites in intensive care. J Hosp Infect. 2017; 95 (1):76–80.

11. Shams AM, Rose □, Edwards JR, Cali S, Harris AD, Jacob JT, et al. Assessment of the Overall and Multidrug-Resistant Organism Bioburden on Environmental Surfaces in Healthcare Facilities. Infect Control Hosp Epidemiol. 2016; 37 (12):1426–32.

12. Silva AS, Deuschle RAN, Garlet CCM. Pesquisa de *Staphylococcus aureus* nas maçanetas das portas dos quartos de um hospital na região Noroeste, Rio Grande do Sul Rev Saúde (Santa Maria). 2012; 38 (1):115–24.

13. Oliveira AC, Damasceno QS. Superfícies do ambiente hospitalar como possíveis reservatórios de bactérias resistentes: uma revisão. Rev Esc Enferm. 2010; 44 (4): 1118–23.

14. Caetano JA, Lima MA, Miranda MC, Serufo JC, Ponte PRL. Identificação de contaminação bacteriana no sabão líquido de uso hospitalar. Rev Esc Enferm. 2011; 45 (1):153–60.

15. Costa DM, Johani K, Melo DS, Lopes LK, Lima LKOL, Tipple AFV, et al. Biofilm contamination of high-touched surfaces in intensive care units: epidemiology and potential impacts. Lett Appl Microbiol. 2019; 68 (4): 267–8.

16. Galvin S, Dolan A, Cahill O, Daniels S, Humphreys H. Microbial monitoring of the hospital environment: why and how? J Hosp Infect. 2012; 82 (3):143–51.

17. Sexton T, Clark P, O’Neill E, Dillane T, Humphreys H. Environmental reservoirs of methicillin-resistant *Staphylococcus aureus* in isolation rooms: correlation with patient isolates and implications for hospital hygiene. J Hosp Infect. 2006; 62 (2):187–94.

18. Dancer CJ. Importance of the environment in methicillin resistant *Staphylococcus aureus* acquisition: the case for hospital cleaning. Lancet Infect Dis. 2008; 8 (2):101–13.

19. Hayden MK, Blom DW, Lyle EA, Moore CG, Weistein RA. Risk of hand or glove contamination after contact with vancomycin resistant Enterococcus or the colonized patients environment. Infect Control Hosp Epidemiol. 2008; 29 (2):149–54.

20. Drees M, Snydman DR, Schmid CH, Barefoot L, Hansjosten K, Vue PM, et al. Prior environmental contamination increases the risk of acquisition of vancomycin-resistant enterococci. Clin Infect Dis. 2008; 46 (5):678–85.

21. Koneman EW, Allen SD, Janda WM, Schreckenberger PC, Winn WC. Diagnóstico microbiológico: texto e atlas colorido. 8. ed. Rio de Janeiro: Guanabara Koogan; 2010, 1565 p.

22. Clinical and Laboratory Standards Institute (CLSI). M100-S24 Performance Standards for Antimicrobial Susceptibility Testing; Twenty-Fourth Informational Supplement. January 2014.

23. Siegel JD, Rhinehart E, Jackson M, Chiarello L. Management of multidrug-resistant organisms in health care settings, 2006. Am J Infect Control. 2007; 35(10 Suppl 2): S165–93.

24. Hidron AI, Edwards JR, Patel J, Horan TC, Sievert DM, Pollock DA. NHSN annual update: Antimicrobial-resistant pathogens associated with healthcare-associated infections: annual summary of data reported to the National Healthcare Safety Network at the Centers for Disease Control and Prevention, 2006–2007. Infect Control Hosp Epidemiol. 2008; 29 (11): 996–1011.

25. Esteves DC, Pereira VC, Souza JM, Keller R, Simões RD, Winkelstroter Eller LK, et al. Influence of biological fluids in bacterial viability on different hospital surfaces and fomites. Am J Infect Control. 2016; 44 (3): 311–4.

26. Moraes CL, Ribeiro NFG, Costa DM, Furlan VG, Palos MAP, Vasconcelos LSNOL. Contaminação de equipamentos e superfícies de unidades de terapia intensiva de uma maternidade pública por *Staphylococcus* coagulase negativa. Rev Patol Trop. 2013; 42 (4): 387–94.

27. Johani K, Abualsaudc D, Costa DM, Hua H, Whiteleye G, Deva A, et al. Characterization of microbial community composition, antimicrobial resistance and biofilm on intensive care surfaces. J Infect Public Health. 2018; 11: 418–24.

28. Carling PC, Parry MF, von Beheren SM. Healthcare Environ-mental Hygiene Study Group. Identifying opportunities to enhance environmental cleaning in 23 acute care hospitals. Infect Control Hosp Epidemiol. 2008; 29 (17): 1–7.

29. Goodman ER, Platt R, Bass R, Onderdonk AB, Yokoe DS, Huang SS. Impact of environmental cleaning intervention on the presence of methicillin-resistant Staphylococcus aureus and vancomycin-resistant enterococci on surfaces in intensive care unit rooms. Infect Control Hosp Epidemiol. 2008; 29: 593–9.

30. Rutala WA, Weber DJ. Healthcare Infection Control Practices Advisory Committee (HICPAC). Guideline for Disinfection and Sterilization in Healthcare Facilities, 2008. Department of health and human services – USA. Centers for diseases control; 2017 Feb. 161p.

31. Lessa FC, Gould CV, McDonald LC. Current status of *Clostridium difficile* infection epidemiology. Clin Infect Dis. 2012; 55 Suppl 2: S65–70.

32. Cortegiani A, Russotto V, Maggiore A, Attanasio M, Naro AR, Raineri SM, et al. Antifungal agents for preventing fungal infections in non-neutropenic critically ill patients. Cochrane Database of Systematic Reviews. 2016; Issue 1. Art. No.: CD004920.

33. Ledwoch K, Dancer SJ, Otter JA, Kerr K, Roposte D, Rushton L, et al. Beware biofilm! Dry biofilms containing bacterial pathogens on multiple healthcare surfaces; a multi-centre study. J Hosp Infect. 2018; 100 (3): 47–56.

34. Nkuwi EJ, Kabanangi F, Joachim A, Rugarabamu S, Majigo M. Methicillin-resistant *Staphylococcus aureus* contamination and distribution in patient’s care environment at Muhimbili National Hospital, Dar es Salaam-Tanzania. BMC Res Notes. 2018; 11(1): 484.

35. Heyba M, Ismaiel M, Alotaibi A, Mahmoud M, Baqer H, Safar A, et al. Microbiological contamination of mobile phones of clinicians in intensive care units and neonatal care units in public hospitals in Kuwait. BMC Infect Dis. 2015; 15: 434–43.

36. Sgro M, Shah PS, Campbell D, Tenuta A, Shivananda S, Lee SK. Early-onset neonatal sepsis: rate and organism pattern between 2003 and 2008. J Perinatol. 2011; 31(12):794–8.

37. Edwards RK, Jamie WE, Sterner D, Gentry S, Counts K, Duff P. Intrapartum antibiotic prophylaxis and early-onset neonatal sepsis patterns. Infect Dis Obstet Gynecol. 2003;11(4): 221–6.

38. Pereira CA, Marra AR, Camargo LF, Pignatari AC, Sukiennik T, Behar PR, et al. Nosocomial Bloodstream Infections in Brazilian Pediatric Patients: Microbiology, Epidemiology, and Clinical Features. PLoS One. 2013; 8 (7): e68144

39. Campos GB, Souza SG, Lobão TN, Silva DCC, Sousa DS, Oliveira OS, et al. Isolation, molecular characteristics and disinfection of methicillin-resistant *Staphylococcus aureus* from ICU units in Brazil. New Microbiol. 2012; (35):183–90.

40. Tajeddina E, Rashidana M, Razaghi M, Javadi SSS, Sherafat SJ, Alebouyeha M, et al. The role of the intensive care unit environment and health-care workers in the transmission of bacteria associated with hospital acquired infections. J Infect Public Health. 2016; 9: 13–23.

41. Rosenthal VD, Bijje H, Maki DG, Mehta Y, Apisarnthanarak A, Medeiros EA, et al. International Nosocomial Infection Control Consortium (INICC) report, data summary of 36 countries, for 2004-2009. Am J of Infect Control. 2012; 40 (5): 396–407.

42. Laabei M, Recker M, Rudkin JK, Aldeljawi M, Gulay Z, Sloan TJ, et al. Predicting the virulence of MRSA from its genome sequence. Genome Res. 2014; 24 (5): 839–49.

43. Heilmann C. Adhesion mechanisms of staphylococci. Adv Exp Med Biol. 2011; 715: 105–23.

44. Judge C, Galvin S, Burke L, Thomas T, Humphreys H, Fitzgerald-Hughes D. Search and you will find: detecting extended-spectrum b-lactamase–producing *Klebsiella pneumoniae* from a patient’s im-mediate environment. Infect Control Hosp Epidemiol. 2013; 34: 534–6.

45. Abreu PM, Farias PG, Gabriel Silva Paiva GS, Almeida AM, Morais PV. Persistence of microbial communities including *Pseudomonas aeruginosa* in a hospital environment: a potential health hazard. BMC Microbiol. 2014, 14:118–27.

46. Ferrareze MVG, Leopoldo VC, Andrade D, Silva MFI, Haas VJ. *Pseudomonas aeruginosa* multirresistente em unidade de cuidados intensivos: desafios que procedem? Acta Paul Enferm. 2007; 20 (1): 7–11.

47. Oliveira C, Malheiros PS, Montagner M, Rossi EM, Brandelli A. Perfil de resistência a antimicrobianos de cepas *Pseudomonas aeruginosa* isoladas de ralos e pios de enfermarias hospitalares em Santa Catarina, Brasil. RBAC. 2011; 43 (3): 192–6.

48. Vickery K, Deva A, Jacombs A, Allan J, Valente P, Gosbell I. Presence of biofilm containing viable multiresistant organisms despite terminal cleaning on clinical surfaces in an intensive care unit. J Hosp Infect. 2012; 80 (1):52–5.

